# MYCN-induced metabolic rewiring creates novel therapeutic vulnerabilities in neuroblastoma

**DOI:** 10.1101/423756

**Authors:** Britta Tjaden, Katharina Baum, Viktoria Marquardt, Mareike Simon, Marija Trajkovic-Arsic, Theresa Kouril, Bettina Siebers, Jan Lisec, Jens T. Siveke, Johannes H. Schulte, Uwe Benary, Marc Remke, Jana Wolf, Alexander Schramm

**Affiliations:** Department of Medical Oncology, West German Cancer Center, University Hospital Essen, University of Duisburg-Essen, Essen, Germany; Mathematical Modeling of Cellular Processes, Max-Delbrück-Center for Molecular Medicine, Berlin, Germany; Division of Pediatric Neuro-Oncogenomics, German Cancer Consortium (DKTK) and German Cancer Research Center (DKFZ), Moorenstraße 5, 40225 Düsseldorf, Germany.; Division of Solid Tumor Translational Oncology, West German Cancer Center, University Hospital Essen, Essen, Germany and German Cancer Consortium (DKTK, partner site Essen) Germany; Molecular Enzyme Technology and Biochemistry (MEB), Biofilm Centre, Faculty of Chemistry, University of Duisburg-Essen, Duisburg, Germany.; Federal Institute for Materials Research and Testing (BAM), Analytical Chemistry, Berlin, Germany; Charité - Universitätsmedizin Berlin, Corporate Member of Freie Universität Berlin, Humboldt-Universität zu Berlin; Berlin Institute of Health (BIH), Berlin, Germany; Department of Pediatric Hematology/Oncology/Stem Cell Transplantation, Berlin, Germany; German Cancer Consortium (DKTK), Heidelberg, Germany; German Cancer Research Center (DKFZ), Heidelberg, Germany.

## Abstract

MYCN is a transcription factor that is aberrantly expressed in many tumor types and is often correlated with poor patient prognosis. Recently, several lines of evidence pointed to the fact that oncogenic activation of MYC family proteins is concomitant with reprogramming of tumor cells to cope with an enhanced need for metabolites during cell growth. These adaptions are driven by the ability of MYC proteins to act as transcriptional amplifiers in a tissue-of-origin specific manner. Here, we describe the effects of MYCN overexpression on metabolic reprogramming in neuroblastoma cells. Ectopic expression of MYCN induced a glycolytic switch that was concomitant with enhanced sensitivity towards 2-deoxyglucose, an inhibitor of glycolysis. Moreover, global metabolic profiling revealed extensive alterations in the cellular metabolome resulting from overexpression of MYCN. Limited supply with either of the two main carbon sources, glucose or glutamine, resulted in distinct shifts in steady-state metabolite levels and significant changes in glutathione metabolism. Interestingly, interference with glutamine-glutamate conversion preferentially blocked proliferation of MYCN overexpressing cells, when glutamine levels were reduced. Thus, our study uncovered MYCN induction and nutrient levels as important metabolic master switches in neuroblastoma cells and identified critical nodes that restrict tumor cell proliferation.

## Introduction

MYC proteins are encoded by a small family of proto-oncogenes consisting of *c*-*MYC*, *N*-*MYC* and *L*-*MYC*. In normal tissues, their expression is tightly regulated, while deregulation of MYC family members has been identified as a driving force in different cancer types. Since specific binding motifs, termed E-boxes, had been identified early on, MYC proteins were considered to be gene-specific transcription factors. This concept has been recently extended by different studies suggesting that deregulated MYC in tumors functions as a general transcriptional amplifier ^1,2,3^. However, MYC(N)-induced and tumor-specific mechanisms of target gene control on transcriptional level have only recently been addressed mechanistically ^4^. The picture emerges that, at least in settings with vastly elevated MYC-levels, enhancer invasion by MYC(N) and associated proteins contributes to tumor-specific MYC/MYCN signatures. Moreover, the concept of MYC-mediated cell autonomous effects to boost tumor cell proliferation has been extended to include restriction of host immune reactions towards a tumor ^5^. Although these efforts led to a better understanding of cell autonomous and cell non-autonomous regulatory circuits governed by oncogenic MYC(N) functions, insights into mechanistic effects on the level of metabolic circuits is still largely lacking.

Deregulated MYC activity comes along with enhanced metabolic stress and increased sensitivity towards apoptosis due to a dependency on continuous supply with nutrients. Glutamine has been identified as a limiting factor for c-MYC dependent cell growth and glutamine deprivation was preferentially inducing apoptosis in MYC-high cells ^6^. In neuroblastoma, the most common solid tumor of childhood, elevated MYCN levels are often found due to amplification of the coding gene, *N*-*Myc*, which is correlated with poor overall survival of affected patients. MYCN overexpression in the absence of MYCN amplification is not prognostic, pointing to the importance of additional genetic factors such as telomerase maintenance for determining disease outcome ^7^. However, forced expression of MYCN is sufficient to induce neuroblastoma in different model organisms including mice ^8,9^ and zebrafish ^10,11^ indicating a causative role for MYCN expression in disease onset and maintenance. Ectopic MYCN expression in neuroblastoma cells is accompanied with increased aggressiveness, but also a higher sensitivity towards drug-induced apoptosis in vivo and in vitro ^12,13^. Interestingly, glutamine deprivation was reported to preferentially sensitize MYCN-amplified cells towards programmed cell death ^14^. MYCN also promoted expression of genes involved in glycolysis including Lactate Dehydrogenase (LDH) and Hexokinase 2 (HK2) in neuroblastoma in vitro ^15^ and in vivo ^16^. The notion that depletion of LDH inhibited MYCN-dependent tumorigenesis and that MYCN cooperated with HIF1α ^15^ suggested that MYCN could also contribute to the Warburg effect. Thus, tumor-specific MYC deregulation directly affects tumor metabolism in multiple pathways.

At present, it is not clear how metabolic dependencies of tumors can be best exploited for therapeutic purposes. It is evident that metabolic features are remarkably flexible, especially those driven by MYC proteins. Moreover, tumor-type specific metabolic adaptations have been identified in MYC-driven tumors, exemplified by the MYC-mediated de novo synthesis of glutamine ^17^. By contrast, MYC-driven liver tumors rather consume glutamine by a process termed glutaminolysis, which allows for fueling into the TCA cycle at the level of α-ketoglutarate by activation of glutaminase, another MYC-target ^18^. MYCN-driven glutaminolysis has been recently suggested as a strategy to treat MYC-driven cancers ^19^. Still, tumor cell-intrinsic response patterns as a consequence of MYCN activation under varying nutrient conditions largely remain to be identified. We thus set out to profile metabolic shifts in neuroblastoma cell lines with inducible MYCN expression and correlate their phenotypic responses upon variations in the two most common carbon sources, glucose and glutamine.

## Materials and methods

### Cell culture and reagents

Neuroblastoma cell lines SHEP, SH-SY5Y, SK-N-AS and SK-N-SH were cultivated in RPMI1640 medium containing 10% fetal bovine serum (FBS) and antibiotics as described ^20-22^. Protocols for generating inducible expression of a gene of interest have been described before ^23^. In brief, cell lines were sequentially transfected with pcDNA6/TR, harboring the tetracycline repressor gene, and pT-REx-DEST30 (ThermoFisher/ Invitrogen) containing *N*-*Myc* cDNA. Single cell clones were selected by limiting dilution in medium containing blasticidine and G418 (ThermoFisher/ Invitrogen). The term “-TR-MYCN” was added to label transfected cells. *N*-*myc* induction was realized by adding 1 μg tetracycline per ml medium. Cell lines were authenticated by STR genotyping prior and post transfections. All reagents used for cell culture were obtained from Gibco/ ThermoFisher. Absence of Mycoplasma sp. in cultivated cells was confirmed by performing PCR with Mycoplasma-specific primers (IDT).

### Metabolic activity and cell viability assays

Metabolic activity was assessed using the 3-(4,5-dimethylthiazol-2-yl)-2,5-diphenyl tetrazolium bromide (MTT) assay. For monitoring cell viability upon inhibition of glutamine metabolism, automated cell handling in a 384 well format was used. Cell Titer Glo assays were performed using a Spark 10M microplate reader (Tecan) as described ^24^.

### Protein analyses and antibodies

Either whole-cell protein extracts or fractionated cell extracts were separated on gradient (4-12%) SDS-PAGE and subsequently used for western blotting. Immunoblot analysis was performed using primary antibodies targeting Hexokinase II (1:500 dilution; #2106); hexokinase I (1:1000; #2804); MYCN (1:1000; #9405); c-Myc (1:1000; #9402); PHGDH (1:1000; #13428, all from Cell Signaling Technology). To ensure equivalent protein loading, blots were stained with a primary antibody raised against β-actin (1:2000 dilution; #A5441, Sigma Aldrich Merck, Munich, Germany). Detection and visualization was achieved using HRP-conjugated goat anti-mouse or anti-rabbit secondary antibodies (GE Healthcare) and the ECL^™^ Prime Western Blotting Detection Reagent (GE Healthcare, #RPN2236), respectively. Data were analyzed using a FusionFX7 detection device (Vilber Lourmat).

### Real-time reverse transcriptase-PCR

Total RNA was isolated from cells using the High Pure RNA Isolation Kit (Roche, #11828665001), and cDNA synthesis was performed using the Transcriptor First Strand cDNA Synthesis Kit (Roche, #04379012001). Expression of target genes was monitored using a StepOnePlus^™^ Real-Time PCR System (Applied Biosystems) and Fast SYBR^®^ Green Master Mix (Roche, #04913914001). Expression values were normalized to β-actin expression and the fold change was calculated using the ΔΔ-Ct method.

### Glycolysis flux assays

Glycolysis flux assays were performed in a Seahorse 96-well XF Cell Culture Microplate using a XF96 sensor cartridge according to the manufacturer’s instructions (Seahorse Bioscience,#102416-100). Briefly, 20,000 cells per well were allowed to grow overnight in a 37°C, 5% CO2 incubator in pyruvate-free RPMI1640 (Gibco/ThermoFisher Scientific, #21875-034, #31870-025, #11879-020) with varying concentrations of L-glutamine (Gibco/ThermoFisher Scientific, #25030-024) and D-glucose (Gibco/ThermoFisher Scientific, #A2494001), 10% fetal bovine serum (Gibco/ThermoFisher Scientific #10270-106) and antiobiotics. On the next day, medium was replaced with pyruvate- and glucose-free Seahorse XF Base Medium (#102353-100) supplemented with 2 mM glutamine according to the manufacturer’s protocol. The microplates were incubated at 37°C for 45 min and then transferred to the microplate stage of a Seahorse XFe96 Extracellular Flux Analyzer to quantify the extracellular acidification rate (ECAR, in [mpH/min]) and the oxygen consumption rate (OCR, in [pmol/min]) using glucose concentrations as indicated. For each step, three separate ECAR and OCR readings using a mix and measure cycling protocol (3 min each) were recorded. Data for each experimental condition were calculated from a minimum of 10 technical replicates.

### Metabolic profiling

SHEP-TR-MYCN cells with and without induction of MYCN were incubated under varying glucose or glutamine concentrations. Upon harvesting, samples were prepared using the automated MicroLab STAR^®^ system (Hamilton). To recover chemically diverse metabolites, proteins were precipitated with methanol under vigorous shaking for 2 min (Glen Mills GenoGrinder 2000) followed by centrifugation. The resulting extract was analyzed either by separate reverse phase (RP)/UPLC-MS/MS with positive ion mode electrospray ionization (ESI), RP/UPLC-MS/MS with negative ion mode ESI or HILIC/UPLC-MS/MS with negative ion mode ESI. The sample extracts were stored overnight under nitrogen before preparation for analysis. All methods utilized a Waters ACQUITY ultra-performance liquid chromatography (UPLC) and a Thermo Scientific Q-Exactive high resolution/accurate mass spectrometer interfaced with a heated electrospray ionization (HESI-II) source and Orbitrap mass analyzer operated at 35,000 mass resolution. Raw data was extracted, peak-identified and QC processed using hardware and software developed by Metabolon Inc..

### Statistical analysis

Statistical analyses of metabolic profiling data were performed using R, version 3.3.2 (available at cran.r-project.org). Prior to analyses, samples were normalized according to protein content determined by Coomassie G-250 based assays. For detailed procedures see Supplemental Methods.

## Results

### Metabolic reprogramming of SHEP cells depends on carbon source supply and MYCN expression

We had previously described a correlation of MYCN with HK2 mRNA expression in SHEP human neuroblastoma cells engineered to stably express MYCN or to activate MYCN only in the presence of tamoxifen ^25^. To better understand the cellular and the metabolic response towards MYCN induction and varying glucose /glutamine ratios, we here used SHEP cells with tightly controlled conditional upregulation of MYCN in the presence of tetracycline (SHEP-TR-MYCN). Expression analyses confirmed both, MYCN induction and concomitant HK2 upregulation in these cells (Fig. 1A). SHEP-TR-MYCN cells were then cultivated with or without MYCN induction and varying glucose and glutamine levels. Of note, MYCN and HK2 induction appeared to be independent of glucose and glutamine concentrations in the medium (Fig. 1A). We defined ten experimental groups that differed in MYCN expression levels and nutrient supply (Table 1) to mimic different carbon source availability in the presence of absence of MYCN. Metabolomic profiling by ultrahigh performance liquid chromatography-tandem mass spectrometry (UPLC-MS/MS) identified 499 compounds, of which 495 (99.2%) were present in at least one third of all samples. Upon normalization to protein content and imputation of missing values, principal component analysis (PCA) demonstrated near perfect separation of samples according to MYCN expression. PCA also separately grouped samples cultivated at low or high concentrations of glucose or glutamine (Fig. 1B). We performed two condition-specific analyses of variance (ANOVAs) for each metabolite, one for constant glutamine conditions to examine the effect of glucose variation, and one for constant glucose conditions to examine the effect of glutamine variation. For each condition, we only analyzed metabolites which were present in at least one third samples considered (Table 1). Induction of MYCN was associated with consistent metabolic shifts, as 144 / 496 (29%, for constant glutamine) or 365 / 486 (75%, for constant glucose) of the metabolites were significantly altered (FDR<0.05) between MYCN high and MYCN low expressing cells (Fig. 1 C-D; Supplemental Figs. 1-2). A comparable number of metabolites were significantly affected when glucose (199 / 496, 40%) or glutamine levels (352 / 486, 73%) were varied. We then examined a possible interaction effect between MYCN expression levels and glucose or glutamine supply. ANOVAs revealed a significant interaction between varying MYCN expression and glucose concentrations affecting 24% of all metabolites (120 / 496 metabolites, FDR < 0.05) when only samples with constant glutamine concentrations were considered (Fig. 1C). An even higher number of metabolites (240 / 486, FDR < 0.05) had a significant interaction effect between MYCN and glutamine when analyzing samples that had constant glucose concentrations (Fig. 1D). Classification of these metabolites according to “super pathways” ^26^ was performed to uncover patterns according to metabolic categories. Here, metabolites classified in the category “energy metabolism” had the highest relative enrichment (Supplemental Figs. 1-2). Thus, global profiling suggested tight interaction of MYCN and carbon source levels in metabolic reprogramming of SHEP cells, while HK2 was confirmed as a MYCN target independent of carbon source availability.

**Figure 1.**
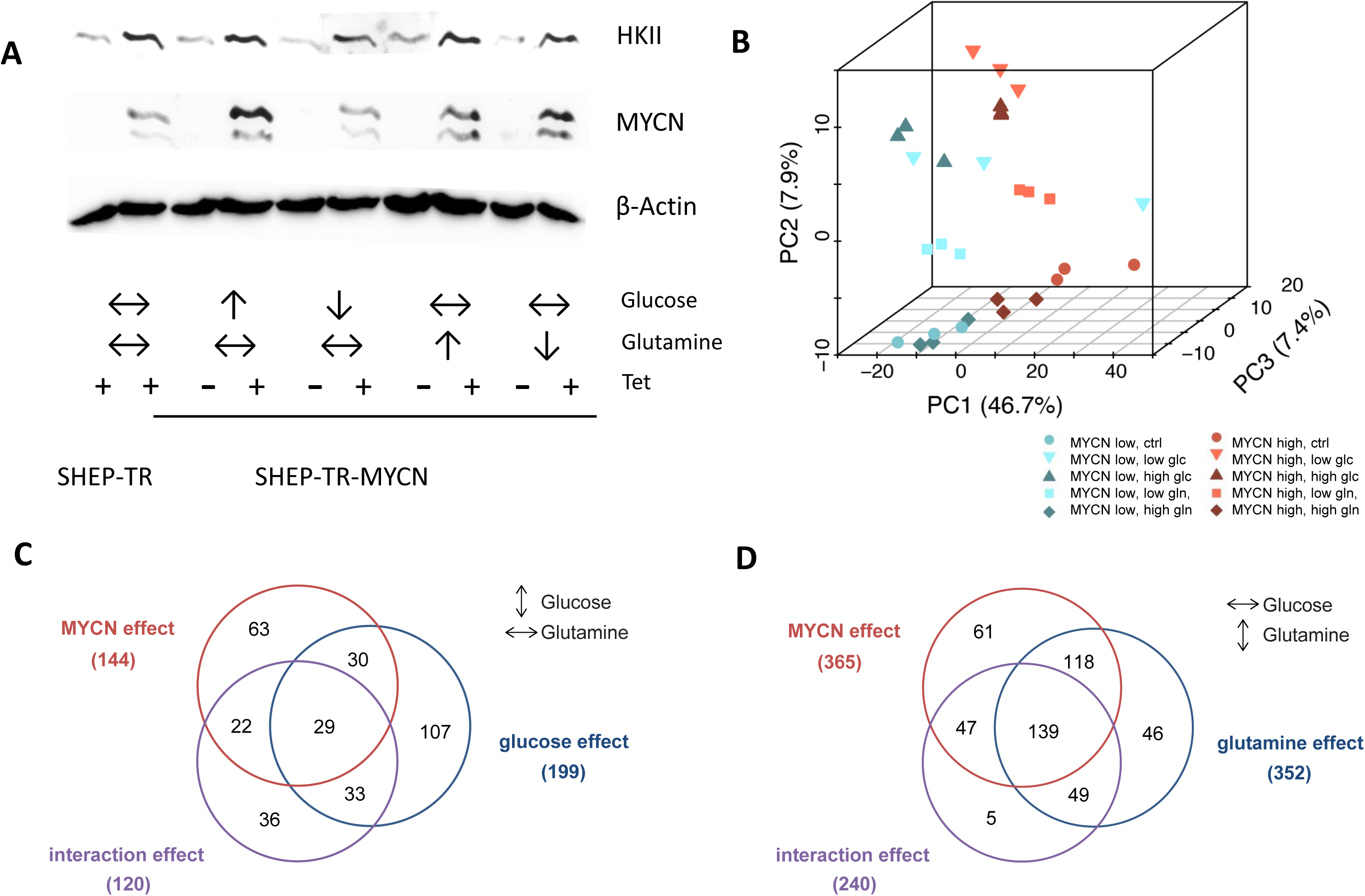
A: Western Blot analyses of Hexokinase II (HKII) and MYCN expression in SHEP-TR-MYCN cells. “+” and “-” refer to induction of MYCN expression by tetracycline (Tet). Arrows indicate high or low concentration of glucose and glutamine, respectively. Double arrows indicate concentrations of both carbon sources used for normal cell culture (cf. Table 1).
B: Principal component analysis (PCA) of metabolic profiling data using SHEP-TR-MYCN cells. In total, 495 metabolites present in at least 10 measurements were included. Each symbol denotes one sample. Blue symbols indicate conditions without MYCN induction (“MYCN low”), while red symbols refer to samples with high MYCN overexpression upon induction by Tet (“MYCN high”). Low and high levels of glucose and glutamine, respectively, were used as defined in Table 1.
C: Numbers of metabolites significantly affected by MYCN, glucose abundance or with significant interaction effect according to bifactorial ANOVAs using sample groups with constant glutamine conditions and varying glucose levels (cf. Table 1).
D: Numbers of metabolites significantly affected by MYCN, glutamine abundance or with significant interaction effect according to bifactorial ANOVAs using the sample groups with constant glucose conditions and varying glutamine levels (cf. Table 1).

**Table 1.**
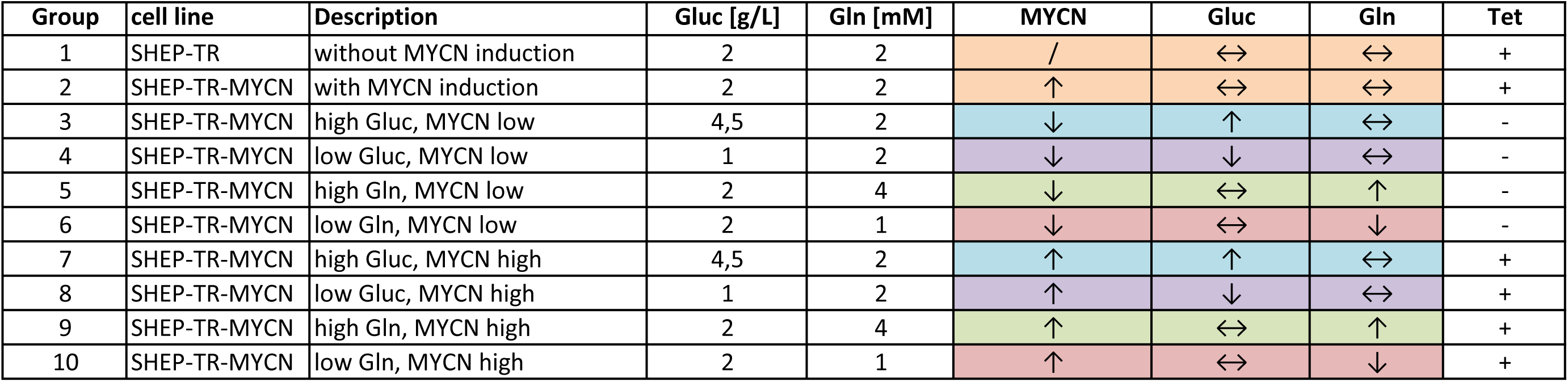
Definition of groups for analyses of metabolite levels in SHEP-TR-MYCN cells with varying levels of carbon source supply as indicated (“Gluc”: glucose; “Gln”: glutamine). MYCN expression in these cells can be induced by addition of tetracycline (“Tet”). Additionally, definition of carbon source concentrations as “high”, “low” or unchanged (indicated by a double arrow) is explained here.

| Group | cell line | Description | Gluc [g/L] | Gln [mM] | MYCN | Gluc | Gln | Tet |
| --- | --- | --- | --- | --- | --- | --- | --- | --- |
| 1 | SHEP-TR | without MYCN induction | 2 | 2 | / | ↔ | ↔ | + |
| 2 | SHEP-TR-MYCN | with MYCN induction | 2 | 2 | ↑ | ↔ | ↔ | + |
| 3 | SHEP-TR-MYCN | high Gluc, MYCN low | 4,5 | 2 | ↓ | ↑ | ↔ | - |
| 4 | SHEP-TR-MYCN | low Gluc, MYCN low | 1 | 2 | ↓ | ↓ | ↔ | - |
| 5 | SHEP-TR-MYCN | high Gln, MYCN low | 2 | 4 | ↓ | ↔ | ↑ | - |
| 6 | SHEP-TR-MYCN | low Gln, MYCN low | 2 | 1 | ↓ | ↔ | ↓ | - |
| 7 | SHEP-TR-MYCN | high Gluc, MYCN high | 4,5 | 2 | ↑ | ↑ | ↔ | + |
| 8 | SHEP-TR-MYCN | low Gluc, MYCN high | 1 | 2 | ↑ | ↓ | ↔ | + |
| 9 | SHEP-TR-MYCN | high Gln, MYCN high | 2 | 4 | ↑ | ↔ | ↑ | + |
| 10 | SHEP-TR-MYCN | low Gln, MYCN high | 2 | 1 | ↑ | ↔ | ↓ | + |

### MYCN shifts the balance from oxidative phosphorylation to glycolysis

To examine if up-regulation of pacemaker enzymes of glycolysis including HK2 were concomitant with a switch in energy harvest, we tested different scenarios in SHEP cells with varying MYCN levels and glucose supply. In all conditions analyzed, we observed a shift towards preferential use of glycolysis upon MYCN induction as seen in higher ECAR levels (Fig. 2A). This effect was independent of cell density (Fig. 2A) and cell number (Fig. 2B). Interestingly, MYCN-induced upregulation of HK2 did not result in altered steady state HK2 enzymatic activity. MYCN low and MYCN high cells also had comparable activity of other glycolytic proteins, glycerinaldehyde-3-phosphate dehydrogenase (GAPDH), lactate dehydrogenase (LDH), triosephosphate isomerase (TIM) and pyruvate kinase (PK, Supplemental Fig. 3). Moreover, subcellular fractionation revealed no difference in HK activity for MYCN low and MYCN high cells in different cellular compartments (Supplemental Fig. 4). While siRNA-mediated knockdown of either HK1 or HK2 was tolerated regardless of MYCN expression levels (Supplemental Fig. 5), blocking of glycolysis downstream of HK by using 2-deoxyglucose preferentially affected SHEP cells with high MYCN levels (Fig. 2C). To check, whether this effect could be recapitulated in other NB cells, we ectopically expressed inducible MYCN in three additional human neuroblastoma cell lines without MYCN gene amplification, SK-N-AS, SK-N-SH and SY5Y. Again, induction of MYCN was accompanied by upregulation of HKII (Fig. 2D). MYCN-mediated sensitization towards 2-DG was also observed for SK-N-AS cells, while SY5Y cells were intrinsically more sensitive to 2-DG regardless of MYCN expression levels (Supplemental Fig. 6, not shown for SY5Y). Thus, MYCN overexpression induced a shift to preferential usage of glycolysis and also increased sensitivity towards inhibition of the glycolytic pathway in a cell line specific manner.

**Figure 2.**
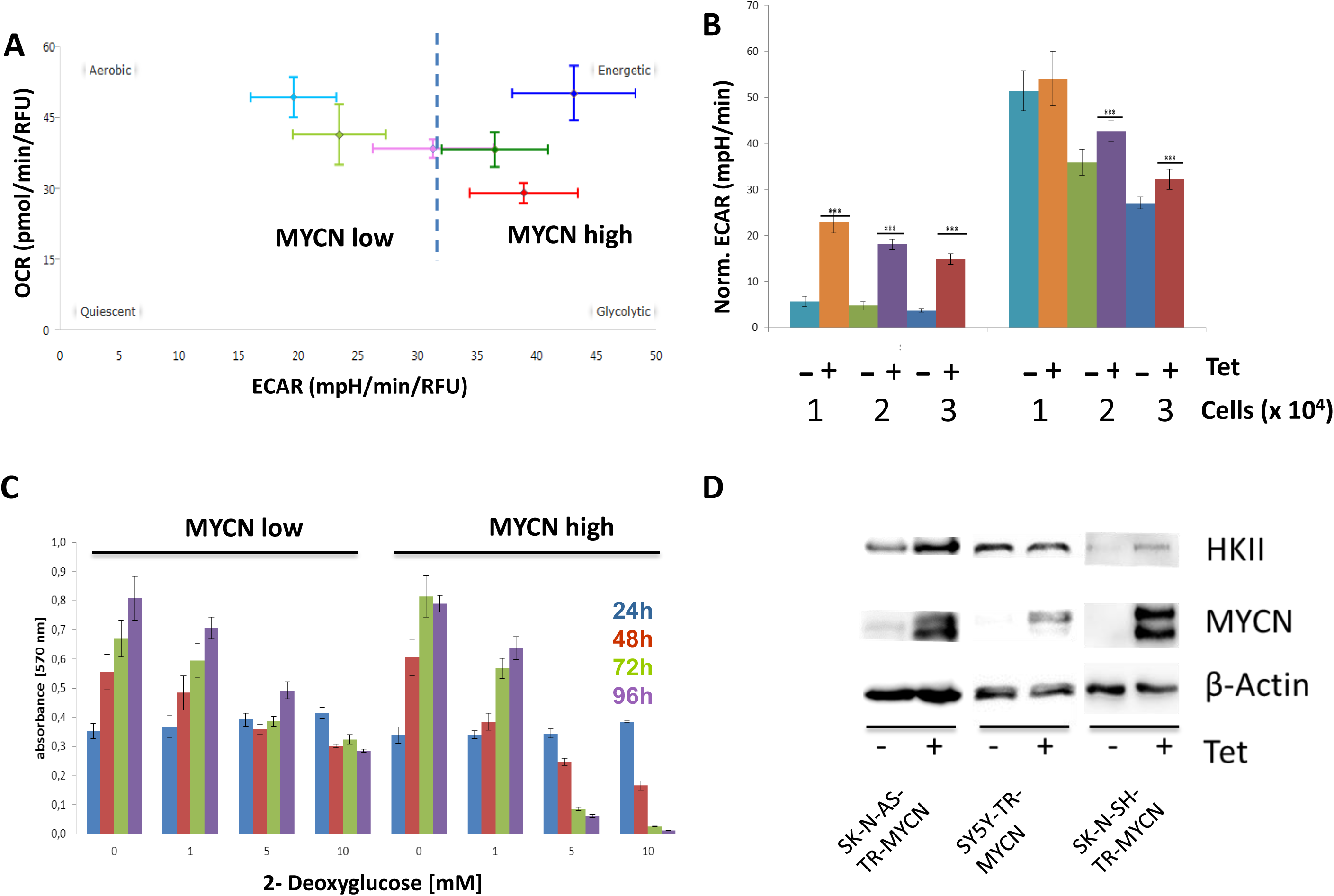
A: Comparison of aerobic respiration and glycolysis indicated by the oxygen consumption rate (OCR) and extracellular acidification rate (ECAR) in SHEP-TR-MYCN cells with (MYCN high) and without induction of MYCN (MYCN low). Colors indicate different glucose (Glc) levels and MYCN expression during cultivation and measurement; MYCN low cells are depicted in pink (1 g Glc /l), light green (2 g Glc /l) and light blue (4.5 g Glc / l), while MYCN high cells are given in red (1 g Glc /l), dark green (2 g Glc /l) and dark blue (4.5 g Glc / l).
B: Glycolysis and glycolytic capacity were monitored by measuring ECAR upon addition of external glucose and oligomycin in SHEP-TR-MYCN cells with and without induction of MYCN by tetracycline using cell numbers as indicated (“Norm. ECAR”: extracellular acidification rate normalized to 10,000 cells).
C: SHEP-TR-MYCN cells were cultivated in the absence (MYCN low) or presence of tetracycline (MYCN high) and incubated with 2-Deoxyglucose. At time points indicated, cell viability was recorded.
D: Hexokinase II (HKII) and MYCN protein expression in SK-N-AS-TR MYCN, SY5Y-TR MYCN and SK-N-SH-TR cells. “+” and “-” refer to addition of tetracycline (Tet), which is regulating MYCN levels.

### MYCN induces cell-type specific responses upon interference with different steps of glutamine biosynthesis

Cellular reprogramming by MYC proteins has been shown to affect glutamine metabolism, consistent with an increased demand of tumor cells for glutamine and metabolic adaptions to fluctuations in nutrient supply (reviewed in ^27^). As for glucose, we observed a high number of significantly altered metabolites depending on MYCN expression levels when glutamine supply was varied. As indicated above, bifactorial ANOVA revealed that 49% of all metabolites displayed a significant interaction effect between MYCN induction and glutamine levels suggesting that these factors are intricately linked to metabolic reprogramming. As MYC proteins upregulate both, the import of glutamine and mitochondrial glutaminolysis, we aimed to investigate the cellular response to inhibition of glutamine metabolism upon MYCN induction. Blocking glutamine uptake by GPNA did not affect cell viability regardless of MYCN expression levels (Fig. 3A). Paradoxically, inhibition of glutaminase by a small molecule inhibitor, CB-839, increased relative cell viability in SK-N-SH TR-MYCN and SHEP-TR-MYCN cells upon MYCN induction when glutamine supply was limited. However, CB-839 did not affect viability of SY5Y TR-MYCN and SK-N-AS TR-MYCN cells over a wide concentration range regardless of MYCN and glutamine levels (Fig. 3B-C and Supplemental Fig. 7). On the other hand, induction of MYCN highly sensitized SY5Y TR-MYCN and SK-N-AS TR-MYCN cells to low doses of the amino acid analogue, DON (6-diazo-5-oxo-L-norleucine), which inhibits the conversion of glutamine to glutamate. SHEP-TR-MYCN cells were resistant to DON concentrations up to 10 μM, while SK-N-SH TR-MYCN cells were sensitized to DON only upon MYCN induction in the presence of low glutamine supply (Fig. 3D-E). Blocking the link between glutaminolysis and the TCA cycle by inhibiting the conversion of glutamate to α-ketoglutarate using aminoxyacetate (AOA) did not affect either of the cell lines regardless of MYCN or glutamine levels (shown exemplarily for SH-EP-TR-MYCN in Fig. 3F). Thus, intervention with glutamine metabolism in neuroblastoma cells uncovered cell-type specific adaptations and metabolic dependencies modulated by MYCN expression and glutamine levels.

**Figure 3.**
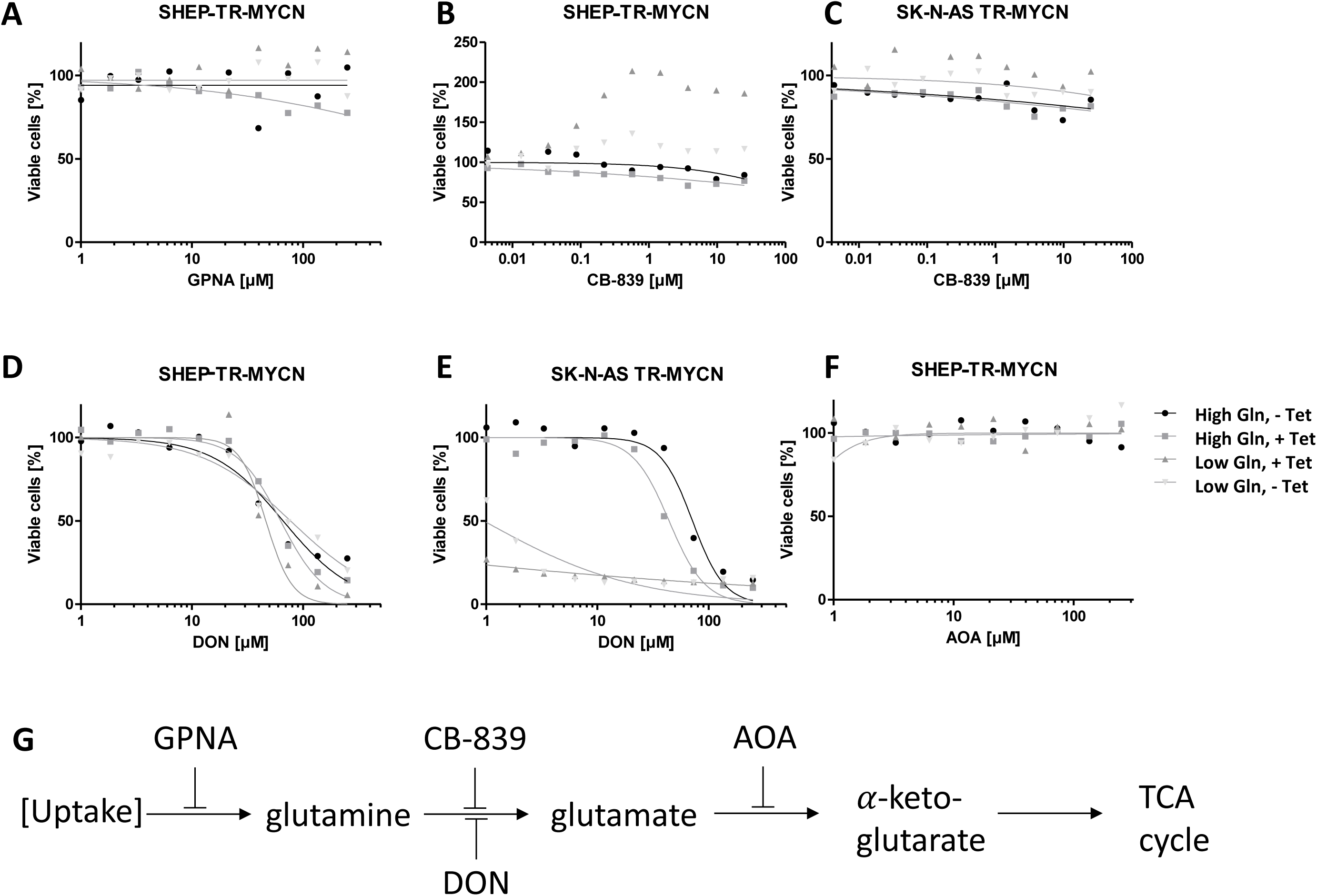
**A-F:** Cell viability depending on small-molecule inhibitors of glutamine metabolism in cell lines indicated. Grey triangles indicate cells incubated in the presence of low Gln with high (dark grey) or low (light grey) MYCN levels; rectangles and circles depict cells cultivated in the presence of high Gln with high (rectangles) or low levels (circles) of MYCN. In experimental conditions indicated, high MYCN expression was induced by addition of tetracycline (“+ Tet”). **G:** Scheme depicting the biochemical steps, in which the small molecule inhibitors of glutamine metabolism are supposed to act (AOA: aminoxyacetate; DON: 6-diazo-5-oxo-L-norleucine).

### MYCN affects the entire pathway of glutathione biosynthesis, but exogenous glutathione does not restore viability of MYCN high cells

In our analysis of metabolome data, classification of known metabolites according to “super pathways” ^26^ was also performed to uncover MYCN induced patterns according to metabolic categories under varying glutamine concentrations. Significant overrepresentation of metabolites upon MYCN induction under these conditions was observed only for the group of “peptides” (Supplemental Fig. 8). Here, 26/26 di-peptides were significantly altered when cells with different MYCN levels were compared. Detailed analyses of this group revealed that 12 of the di-peptides contained gamma-glutamyl residues and were linked to glutathione metabolism. Additional metabolites annotated as members of “glutathione metabolism” ^28^ were also significantly altered by MYCN levels independent of glutamine supply (32/34 metabolites, Fig. 4A); glutathione-associated metabolites were significantly over-represented among the MYCN-affected metabolites (p-value 0.003, Fisher’s exact test). Strikingly, almost all metabolites involved in glutathione metabolism, including glutamine, were found at significantly lower levels in MYCN high cells compared to MYCN low cells regardless of glutamine supply (Fig. 4B). While this suggested a limiting role of glutathione availability for MYCN high cells, DON-mediated reduced viability of SHEP-TR-MYCN cells could not be rescued by exogenous glutathione supply (Fig. 4C). Additionally, increasing reactive oxygen species (ROS) by mono- or dimethyl fumarate (MMF, DMF) did also not affect cell viability upon MYCN induction and varying glutamine supply (Fig. 4D, Supplemental Figure 9). These results indicate that MYCN induction causes a significant depletion of glutamine-related metabolites including glutathione, and that the resulting glutamine addiction cannot be attributed to modulation of ROS levels.

**Figure 4.**
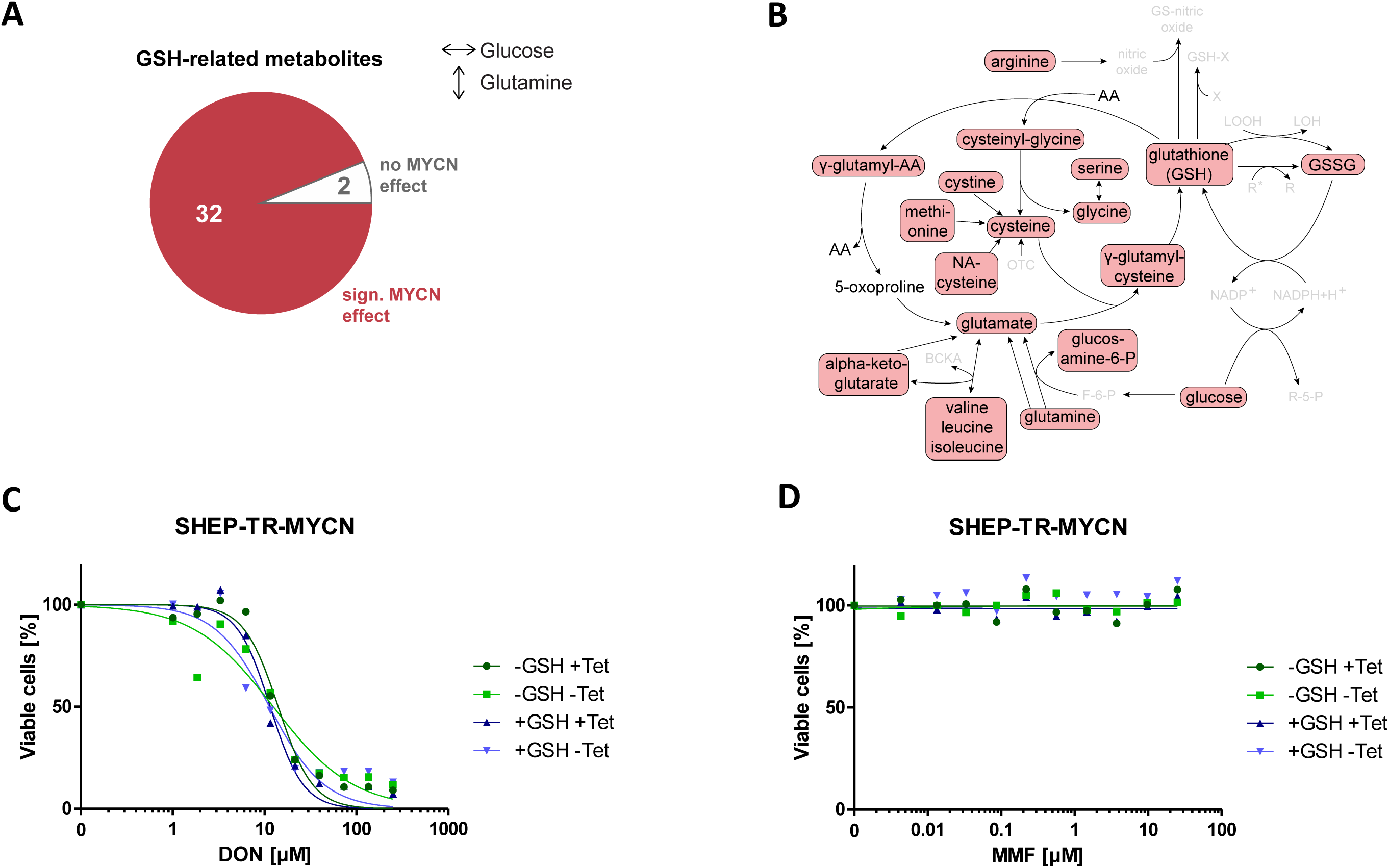
A: Data presented in Fig. 1D were re-analyzed to visualize a significant MYCN-mediated effect on glutathione (GSH) metabolism. In total, 32 out of 34 GSH-related metabolites are significantly affected by MYCN levels according to ANOVA.
B: Main routes of the glutathione pathway (modified from ^28^). Metabolites which are significantly affected by MYCN for the sample groups under constant glucose and varying glutamine concentration in the medium are marked by red boxes. Interestingly, all significantly altered species are found to be decreased upon MYCN induction. Metabolites, that have not been detected by UHPLC are greyed (AA: amino acid, BCKA: branched chain keto acids, F-6-P: fructose-6-phosphate, GSSG: glutathione disulfide, LOOH: lipid hydroxyperoxide, LOH: lipid peroxyl radical, NA-cysteine: N-acetyl-cysteine, OTC: oxothiazolidine-4-carboxylate, R-5-P: ribose-5-phosphate, R, R+: radicals, X: xenobiotic).
C, D: Redox modulation by addition of GSH (100 mM) in the presence of varying concentrations of DON (C) or the ROS-inducer mono-methyl fumarate (MMF, D) in SHEP TR-MYCN cells with or without induction of MYCN (“+Tet” and “−Tet”, respectively).

## Discussion

The wealth of data on plasticity of tumor cells suggests that blocking of single metabolic pathways may not be sufficient for long-term control of cancer growth. However, it is crucial to analyze oncogene-induced adaptations of metabolic pathways on a cellular level to understand mechanisms of metabolic plasticity contributing to development of therapy resistance in cancer. Still, tumor cell-centric assays are limited as they will miss interactions with the tumor microenvironment. A prime example is the notion that K-RAS driven lung tumor cells are glutamine dependent in vitro, as they need glutamine-derived α-ketoglutarate to enter the TCA cycle, but are glutamine-independent in vivo due to fueling of the TCA cycle by glucose ^29^. Expression of glutamine synthetase (GS) by cells of the tumor microenvironment might explain compensatory effects in cancer cells with reduced GS activity ^30^. It has also been argued that routinely applied cell culture conditions will not appropriately mimic metabolic conditions found in human plasma ^31^. Thus, experiments to model tumor cell responses to oncogenic activation in vitro have to be carefully designed. We here investigated the effects of ectopic MYCN expression in neuroblastoma cells lines in the presence of high or low supply with glucose or glutamine, respectively, as an approach to mimic both, oncogene-induced changes in metabolic patterns and response to altered carbon source supply. For metabolic profiling of these conditions, we chose SHEP NB cells, in which we had previously noticed MYCN-induced HK2 upregulation indicating metabolic reprogramming ^16,25^. Using a newly adapted system for inducible MYCN expression, designated SHEP-TR-MYCN, we were able to demonstrate that HK2 induction by MYCN was independent of carbon source supply (Fig. 1A). Moreover, MYCN-dependent upregulation of HK2 could be confirmed in other NB cell lines (Fig. 2D). Metabolome analyses revealed that both, carbon source and MYCN expression levels, cause significant and specific changes allowing for separation of these experimental groups by principal component analyses (Fig. 1B). Of note, we also observed significant interaction effects between MYCN and carbon source levels (Fig. 1C, D) suggesting that MYCN induced reprogramming also affects downstream metabolic processes under varying nutrient conditions. While our analyses uncovered that MYCN-mediated regulation of metabolites affects all pathways investigated, classification of MYCN-regulated metabolites confirmed enrichment for metabolites within the categories “energy metabolism” and a highly significant enrichment for metabolites involved in the glutathione pathway. We thus focused on analyses of two main pathways for energy harvest, glycolysis and glutamine metabolism, as the latter feeds also into glutathione biosynthesis. Indeed, we found a shift towards preferential use of glycolysis in MYCN-induced cells and an increased sensitivity towards glycolysis inhibition (Fig. 2). Glycolysis addiction in cancer cells with high levels of MYC has been described and linked to mTOR pathway activation ^32^. Interestingly, we and others previously noticed an increased sensitivity of MYCN-dependent neuroblastoma towards mTOR inhibition ^25,33^, suggesting that simultaneous blockade of glycolysis and mTOR would be an option to interfere with both, metabolism and survival pathways.

The addiction of MYCN-amplified neuroblastoma cells to glutamine availability has been described ^14^. A recent report indicated higher glutamate / glutamine ratios in neuroblastoma compared to other cell types ^34^. These findings suggested that neuroblastoma can be categorized as a “glutamine-consuming” rather than a “glutamine-producing” tumor type. Of note, c-MYC regulates glutamine metabolism, at least in part, by repressing miR-23a and miR-23b resulting in enhanced expression of their target gene, glutaminase ^18^. Recent reports pointed to a possible feedback mechanism, as MYC translation itself is regulated by intracellular levels of glutamine-derived adenosine nucleotides via a sequence element in the 3’-UTR of the *MYC* mRNA ^6^. In our model cell lines, this effect cannot be analyzed in the absence of endogenous regulator elements in the 3’UTR of the MYCN encoding vectors. The emerging picture, however, is far from clear: blocking glutamine uptake and its metabolic conversion to α-ketoglutarate was well tolerated in all conditions analyzed, indicating compensatory mechanisms. Paradoxical effects were observed for CB-839, an inhibitor of mitochondrial glutaminase (GLS1) that is currently tested in early clinical trials for treatment of several solid tumors multiple myeloma and acute myeloid leukemia ^35^: while two of four NB cell lines did not respond to CB-839, MYCN expression combined with glutamine starvation enhanced metabolic activity of SK-N-SH and SHEP cells. The latter two cell lines were also resistant to DON, which is an irreversible glutamine-competitive inhibitor. However, limiting glutamine concentrations increased sensitivity towards DON in SY5Y and SKNAS cells, while MYCN expression enhanced DON sensitivity 10-fold in SK-N-SH cells. Dissection of cellular responses on the level of the entire metabolome in SHEP cells revealed significant changes in GSH-related metabolites upon variation of glutamine availability, which affected the vast majority of intermediates on the route from glutamine to GSH/GSSG (Fig. 4 A,B). We thus hypothesized that glutamine addiction of MYCN high cells could be attributed to an increased sensitivity towards ROS production. Increased demand for ROS detoxification has been recognized as a vulnerability of cancer cells in the past years. The concept has also helped to understand the mechanisms of action for cytotoxic drugs including As_2_O_3_ and anthracyclines, which are effective at least in part by GSH oxidation and increased ROS production, respectively ^36^. However, sensitivity of MYCN high cells to inhibition of glutamine metabolism could not be rescued by external glutathione supply. Moreover, MYCN high cells were not significantly more sensitive to the ROS-inducing agent, dimethyl fumarate, which was described to suppress neuroblastoma cell proliferation ^19^. Thus, preferential killing of MYCN high cells in the presence of glutamine metabolism inhibitors and limited glutamine supply cannot be explained by increased consumption of glutathione and increased ROS sensitivity.

As mentioned above, strategies aiming to limit carbon source availability of inhibition of primary metabolic pathways have to take into account not only plasticity of tumor cells themselves but also that other cell types in the tumor microenvironment could serve as a carbon source in vivo ^30^. Thus, our results confirm interference with both glucose and glutamine availability as two possible avenues for exploiting metabolic dependencies of MYCN high cells, which have to be further explored in in vivo models.

## Legends to Supplemental Figures

**Supplemental Figure 1:** Metabolomic changes affected by MYCN and media conditions when glucose concentrations were varied. Univariate, bifactorial ANOVA was applied for each of the 496 metabolites present in at least six samples. Numbers of significantly altered metabolites (FDR<0.05 after multiple testing correction) that depend on MYCN expression level (high or low; MYCN effect), medium condition (low glucose, normal glucose, high glucose; glucose effect) or their combined effect (interaction effect) in the 18 samples with varying glucose and constant glutamine abundances are given. Of 496 metabolites, 24% displayed a significant interaction effect (Venn diagram).

Numbers of metabolites with significant interaction effect between MYCN expression and glucose concentration in each category of the “super pathway” classification and their percentage (from the number of metabolites measured in each category) are reported in the lower table.

**Supplemental Figure 2:** Metabolomic changes affected by MYCN and media conditions when glutamine concentrations were varied. Univariate, bifactorial ANOVA was applied for each of the 486 metabolites present in at least six samples. Numbers of significantly altered metabolites depending on MYCN expression level (high or low; MYCN effect), medium condition (low glutamine, normal glutamine, high glutamine; glutamine effect) or their combined effect (interaction effect) in the 18 samples with varying glutamine and constant glucose abundances are given. Of 486 metabolites, 49% displayed a significant interaction effect (Venn diagram). The numbers of metabolites with significant interaction effect between MYCN expression and glutamine concentration in each category of the “super pathway” classification and their percentage (from the number of metabolites measured in each category) are reported in the lower table.

**Supplemental Figure 3:** Specific enzymatic activity in cell extracts from SHEP-TR-MYCN cells for the glycolytic enzymes hexokinase (HK), triosephosphate isomerase (TIM), glycerinealdehyde-3-phosphate dehydrogenase (GAPDH), pyruvate kinase (PK) and lactate dehydrogenase (LDH) was determined as described ^37,38^. Enzymatic activities were determined in “MYCN low” (blue) and “MYCN high” (red) cells, when “MYCN high” was induced by addition of tetracycline.

**Supplemental Figure 4:** Hexokinase (HK) activity was determined in SHEP-TR-MYCN cells under varying conditions as indicated. Left: HK activity in total cell extracts, right: Relative HK activity upon subcellular fractionation of cell extracts. No significant differences were observed between these conditions.

**Supplemental Figure 5:** Left: Isoform-specific knockdown of hexokinase I and II (HKI and HKII) in SHEP-TR-MYCN cells was achieved by specific siRNAs when compared to untransfected (mock) or scrambled RNA (scrRNA, siRNA4 targets only HKI, while siRNA 3 is specific for HKII). Down-regulation of HKI and HKII was independent of MYCN-induction by tetracycline (Tet). Right: Neither downregulation of HKI nor HKII resulted in a significant reduction of cell viability compared to untransfected (mock) or scrambled RNA (scrRNA). Control-transfected SHEP-TR cells (“mock(TR)”) and SHEP-TR-MYCN cells transfected with an siRNA targeting another metabolic enzyme, PHGDH, were used as additional controls.

**Supplemental Figure 6:** SK-N-AS TR-MYCN cells were treated with 2-deoxyglucose for indicated time points (purple: t=0; red: t= 24 h; yellow: t= 48 h; light blue: t= 72 h) and the cell viability was determined.

**Supplemental Figure 7:** Cell viability depending on a small-molecule inhibitors of glutamine metabolism, CB-839, in cell lines indicated. Grey triangles indicate cells incubated in the presence of low Gln with high (dark grey) or low (light grey) MYCN levels; rectangles and circles depict cells cultivated in the presence of high Gln with high (rectangles) or low levels (circles) of MYCN.

**Supplemental Figure 8:** Metabolites being significantly affected by MYCN levels when glutamine concentrations were varied (according to an ANOVA, see also Fig. 1D) were classified according to categories of “super pathways”. Numbers reflect the fraction of the total number of metabolites in each category. Upon correction for multiple testing, only the category “peptides” was found as significantly over-represented (corrected Fisher’s p-value < 0.05).

**Supplemental Figure 9:** Treatment of SHEP-TR-MYCN with mono- or dimethyl fumarate (MMF, DMF) in the presence of high (+Gln) and low (-Gln) glutamine medium concentrations, respectively.

Additional variables were GSH concentration (100 μM= “+GSH”, no external GSH= “-GSH”) and high or low MYCN expression induced by tetracycline (+Tet), which was absent in controls (-Tet).

## References

1. Lorenzin, F. et al. Different promoter affinities account for specificity in MYC-dependent gene regulation. eLife 5 (2016).

2. Walz, S. et al. Activation and repression by oncogenic MYC shape tumour-specific gene expression profiles. Nature 511, 483–487 (2014).

3. Lin, C. Y. et al. Transcriptional amplification in tumor cells with elevated c-Myc. Cell 151, 56–67 (2012).

4. Zeid, R. et al. Enhancer invasion shapes MYCN-dependent transcriptional amplification in neuroblastoma. Nature genetics 50, 515–523 (2018).

5. Kortlever, R. M. et al. Myc Cooperates with Ras by Programming Inflammation and Immune Suppression. Cell 171, 1301–1315.e14 (2017).

6. Dejure, F. R. et al. TheMYCmRNA 3’-UTR couples RNA polymerase II function to glutamine and ribonucleotide levels. The EMBO journal 36, 1854–1868 (2017).

7. Peifer, M. et al. Telomerase activation by genomic rearrangements in high-risk neuroblastoma. Nature 526, 700–704 (2015).

8. Weiss, W. A. Aldape, K. Mohapatra, G. Feuerstein, B. G. & Bishop, J. M. Targeted expression of MYCN causes neuroblastoma in transgenic mice. The EMBO journal 16, 2985–2995 (1997).

9. Althoff, K. et al. A Cre-conditional MYCN-driven neuroblastoma mouse model as an improved tool for preclinical studies. Oncogene 34, 3357–3368 (2015).

10. Zhu, S. et al. LMO1 Synergizes with MYCN to Promote Neuroblastoma Initiation and Metastasis. Cancer cell 32, 310–323.e5 (2017).

11. Zhu, S. et al. Activated ALK collaborates with MYCN in neuroblastoma pathogenesis. Cancer cell 21, 362–373 (2012).

12. Lutz, W. Fulda, S. Jeremias, I. Debatin, K. M. & Schwab, M. MycN and IFNgamma cooperate in apoptosis of human neuroblastoma cells. Oncogene 17, 339–346 (1998).

13. Fulda, S. Lutz, W. Schwab, M. & Debatin, K. M. MycN sensitizes neuroblastoma cells for drug-induced apoptosis. Oncogene 18, 1479–1486 (1999).

14. Qing, G. et al. ATF4 regulates MYC-mediated neuroblastoma cell death upon glutamine deprivation. Cancer cell 22, 631–644 (2012).

15. Qing, G. et al. Combinatorial regulation of neuroblastoma tumor progression by N-Myc and hypoxia inducible factor HIF-1alpha. Cancer research 70, 10351–10361 (2010).

16. Schulte, J. H. et al. Deep sequencing reveals differential expression of microRNAs in favorable versus unfavorable neuroblastoma. Nucleic acids research 38, 5919–5928 (2010).

17. Bott, A. J. et al. Oncogenic Myc Induces Expression of Glutamine Synthetase through Promoter Demethylation. Cell metabolism 22, 1068–1077 (2015).

18. Gao, P. et al. c-Myc suppression of miR-23a/b enhances mitochondrial glutaminase expression and glutamine metabolism. Nature 458, 762–765 (2009).

19. Wang, T. et al. MYCN drives glutaminolysis in neuroblastoma and confers sensitivity to an ROS augmenting agent. Cell death & disease 9, 220 (2018).

20. Ackermann, S. et al. Polo-like kinase 1 is a therapeutic target in high-risk neuroblastoma. Clinical cancer research : an official journal of the American Association for Cancer Research 17, 731–741 (2011).

21. Schulte, J. H. et al. Transcription factor AP2alpha (TFAP2a) regulates differentiation and proliferation of neuroblastoma cells. Cancer letters 271, 56–63 (2008).

22. Schulte, J. H. et al. Microarray analysis reveals differential gene expression patterns and regulation of single target genes contributing to the opposing phenotype of TrkA- and TrkB-expressing neuroblastomas. Oncogene 24, 165–177 (2005).

23. Pajtler, K. W. et al. Neuroblastoma in dialog with its stroma: NTRK1 is a regulator of cellular crosstalk with Schwann cells. Oncotarget 5, 11180–11192 (2014).

24. Stenzel, K. et al. Alkoxyurea-Based Histone Deacetylase Inhibitors Increase Cisplatin Potency in Chemoresistant Cancer Cell Lines. Journal of medicinal chemistry 60, 5334–5348 (2017).

25. Schramm, A. et al. Next-generation RNA sequencing reveals differential expression of MYCN target genes and suggests the mTOR pathway as a promising therapy target in MYCN-amplified neuroblastoma. International journal of cancer 132, E106-15 (2013).

26. Krumsiek, J. et al. Gender-specific pathway differences in the human serum metabolome. Metabolomics : Official journal of the Metabolomic Society 11, 1815–1833 (2015).

27. Dejure, F. R. & Eilers, M. MYC and tumor metabolism. Chicken and egg. The EMBO journal 36, 3409–3420 (2017).

28. Wu, G. Fang, Y.-Z. Yang, S. Lupton, J. R. & Turner, N. D. Glutathione metabolism and its implications for health. The Journal of nutrition 134, 489–492 (2004).

29. Davidson, S. M. et al. Environment Impacts the Metabolic Dependencies of Ras-Driven Non-Small Cell Lung Cancer. Cell metabolism 23, 517–528 (2016).

30. Castegna, A. & Menga, A. Glutamine Synthetase: Localization Dictates Outcome. Genes 9 (2018).

31. Cantor, J. R. et al. Physiologic Medium Rewires Cellular Metabolism and Reveals Uric Acid as an Endogenous Inhibitor of UMP Synthase. Cell 169, 258-272.e17 (2017).

32. Pusapati, R. V. et al. mTORC1-Dependent Metabolic Reprogramming Underlies Escape from Glycolysis Addiction in Cancer Cells. Cancer cell 29, 548–562 (2016).

33. Vaughan, L. et al. Inhibition of mTOR-kinase destabilizes MYCN and is a potential therapy for MYCN-dependent tumors. Oncotarget 7, 57525–57544 (2016).

34. Kohe, S. E. et al. Metabolic profiling of the three neural derived embryonal pediatric tumors retinoblastoma, neuroblastoma and medulloblastoma, identifies distinct metabolic profiles. Oncotarget 9, 11336–11351 (2018).

35. Jacque, N. et al. Targeting glutaminolysis has antileukemic activity in acute myeloid leukemia and synergizes with BCL-2 inhibition. Blood 126, 1346–1356 (2015).

36. Panieri, E. & Santoro, M. M. ROS homeostasis and metabolism. A dangerous liason in cancer cells. Cell death & disease 7, e2253 (2016).

37. Schramm, A. Siebers, B. Tjaden, B. Brinkmann, H. & Hensel, R. Pyruvate kinase of the hyperthermophilic crenarchaeote Thermoproteus tenax: physiological role and phylogenetic aspects. Journal of bacteriology 182, 2001–2009 (2000).

38. Siebers, B. Wendisch, V. F. & Hensel, R. Carbohydrate metabolism in Thermoproteus tenax: in vivo utilization of the non-phosphorylative Entner-Doudoroff pathway and characterization of its first enzyme, glucose dehydrogenase. Archives of microbiology 168, 120–127 (1997).

